# Gene-Specific Predictability of Protein Levels from mRNA Data in Humans

**DOI:** 10.1101/399816

**Authors:** Alief Moulana, Adriana Scanteianu, DeAnalisa Jones, Alan D. Stern, Mehdi Bouhaddou, Marc R. Birtwistle

**Author notes:** To whom all correspondence should be addressed or.

## Abstract

Transcriptomic data are widely available, and the extent to which they are predictive of protein abundances remains debated. Using multiple public databases, we calculate mRNA and mRNA-to-protein ratio variability across human tissues to quantify and classify genes for protein abundance predictability confidence. We propose that such predictability is best understood as a spectrum. A gene-specific, tissue-independent mRNA-to-protein ratio plus mRNA levels explains ∼80% of protein abundance variance for more predictable genes, as compared to ∼55% for less predictable genes. Protein abundance predictability is consistent with independent mRNA and protein data from two disparate cell lines, and mRNA-to-protein ratios estimated from publicly-available databases have predictive power in these independent datasets. Genes with higher predictability are enriched for metabolic function, tissue development/cell differentiation roles, and transmembrane transporter activity. Genes with lower predictability are associated with cell adhesion, motility and organization, the immune system, and the cytoskeleton. Surprisingly, many genes that regulate mRNA-to-protein ratios are constitutively expressed but also exhibit ratio variability, suggesting a general autoregulation mechanism whereby protein expression profile changes can be implemented quickly, or homeostatic sensing stabilizes protein abundances under fluctuating conditions. Gene classifications and their mRNA-to-protein ratios are provided as a resource to facilitate protein abundance predictions by others.

## Introduction

The process of gene expression begins with the transcription of genes to make mRNA, followed by the translation of mRNA to make protein, implying that an increase in transcription rate and higher mRNA levels should produce higher protein levels. However, it is now reasonably well accepted that mRNA and protein levels correlate poorly, particularly for human cells (Liu *et al*, 2016). This conclusion is typically based on log-log scatterplots of mRNA levels versus protein levels from genome-scale measurements, and evaluated quantitatively via the Pearson correlation (*ρ*) or coefficient of determination (*R*^*2*^), which ranges from ∼0.1-0.4 (Maier *et al*, 2009; de Sousa Abreu *et al*, 2009). This lack of correlation has been attributed to regulation of both translation and protein degradation, although the former may have a greater role (Schwanhäusser *et al*, 2011). Translation rate regulation mechanisms include microRNAs (miRNAs) that can inhibit translation initiation (Jonas & Izaurralde, 2015), secondary structure of the mRNA itself (López & Pazos, 2015; Faure *et al*, 2016), RNA binding proteins that can alter affinity for translation factors and/or ribosomes (Gerstberger *et al*, 2014), and protein folding (Cheng *et al*, 2016). Protein degradation rate can be regulated by post-translational modifications, for example by ubiquitination (Ravid & Hochstrasser, 2008), hydroxylation (Fong & Takeda, 2008), or phosphorylation (Swaney *et al*, 2013).

On the other hand, more recent work has supported the notion that with precise enough protein level measurements, the ratio of mRNA-to-protein for a particular gene is found to be well conserved across human tissues and/or cell types and allows very good prediction of protein levels from mRNA levels on a gene-by-gene basis (Wilhelm *et al*, 2014; Edfors *et al*, 2016). Yet, this view has been challenged, in that translational and protein half-life regulation is differentially implemented across cell and tissue types, as well as modulated by environmental signals, and further, that gene-specific metrics of protein prediction reliability may be misleading (Fortelny *et al*, 2017; Franks *et al*, 2017; Perl *et al*, 2017; Parca *et al*, 2018; Wilhelm *et al*, 2017). Therefore, the extent to which protein levels might be predicted from mRNA data remains unclear and an unresolved topic of widespread biological importance.

Here, we leverage publicly-available, human mRNA and protein abundance datasets— from the Human Proteome Map (HPM) (Kim *et al*, 2014), Proteomics DataBase (ProteomicsDB) (Wilhelm *et al*, 2014), and Genotype-Tissue Expression Consortium (GTEx) (Ardlie *et al*, 2015; Melé *et al*, 2015; The GTEx Consortium *et al*, 2015) to parse genes into four categories strongly influencing confidence in protein abundance prediction from mRNA levels using gene-specific mRNA-to-protein ratios. However, we argue that such predictability is best understood as a spectrum. In addition, we find protein abundance predictability to be consistent with mRNA and protein data collected from two distinct cell lines, the U87 glioma and MCF10A breast epithelial lines. Importantly, ratios estimated from ProteomicsDB improve prediction of protein levels in these independent datasets. We find genes with higher predictability to be enriched for metabolic function, tissue development/cell differentiation roles, and transmembrane transporter activity. Genes with lower predictability are associated with cell adhesion, motility and organization, the immune system, and the cytoskeleton. Surprisingly, genes whose transcripts are constitutively expressed but have less predictable protein levels were found to be enriched for post-transcriptional and translational regulation, suggesting a general autoregulation mechanism by which translationally regulated genes regulate themselves. We provide gene mRNA-to-protein ratios across tissues and their predictability classifications in comprehensive supplementary tables.

## Results and Discussion

### Protein Abundance Variability Explained by mRNA Levels and by mRNA-to-Protein Ratio

For a particular gene *i*, a simple linear relationship between the mRNA levels *m*_*i*_ and protein levels *p*_*i*_ (lumping splice isoforms) can be obtained by casting translation and protein degradation as effective first-order processes, and solving for a steady-state condition (de Sousa Abreu *et al*, 2009), yielding

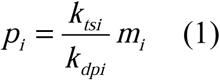

where *k*_*tsi*_ is the translation rate constant [# of protein molecules / # of mRNA molecules / time], and *k*_*dpi*_ is the protein degradation rate constant [1/time]. Often, mRNA and protein levels vary several orders of magnitude across genes, and therefore their log is analyzed instead

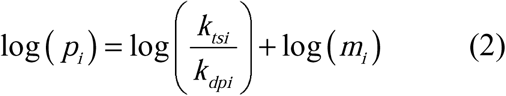

Equations 1 and 2 show that one would expect high linear correlation between mRNA levels and protein levels (or their logs) globally if the rate constant ratio *k*_*tsi*_*/k*_*dpi*_ were relatively invariant across all genes. Given the numerous regulatory mechanisms controlling translation and protein degradation, it is not surprising that this ratio shows significant variability, over an approximately 10,000-fold range in mouse fibroblasts for the 4,247 genes where this ratio was available (Fig. 1A) (Schwanhäusser *et al*, 2011). By sampling this distribution (well fit by a log-normal), along with that of mRNA expression levels (which are also approximately log-normal and uncorrelated with the rate constant ratio—Fig. S1A), we observed simulated mRNA-protein level correlations that yield the unexpectedly low R^2^ values that are typically observed in human cells (Fig. 1B). Thus, because rate constant ratios vary widely across the genome, prediction of protein levels from just mRNA levels will remain poor.

**Figure 1.**
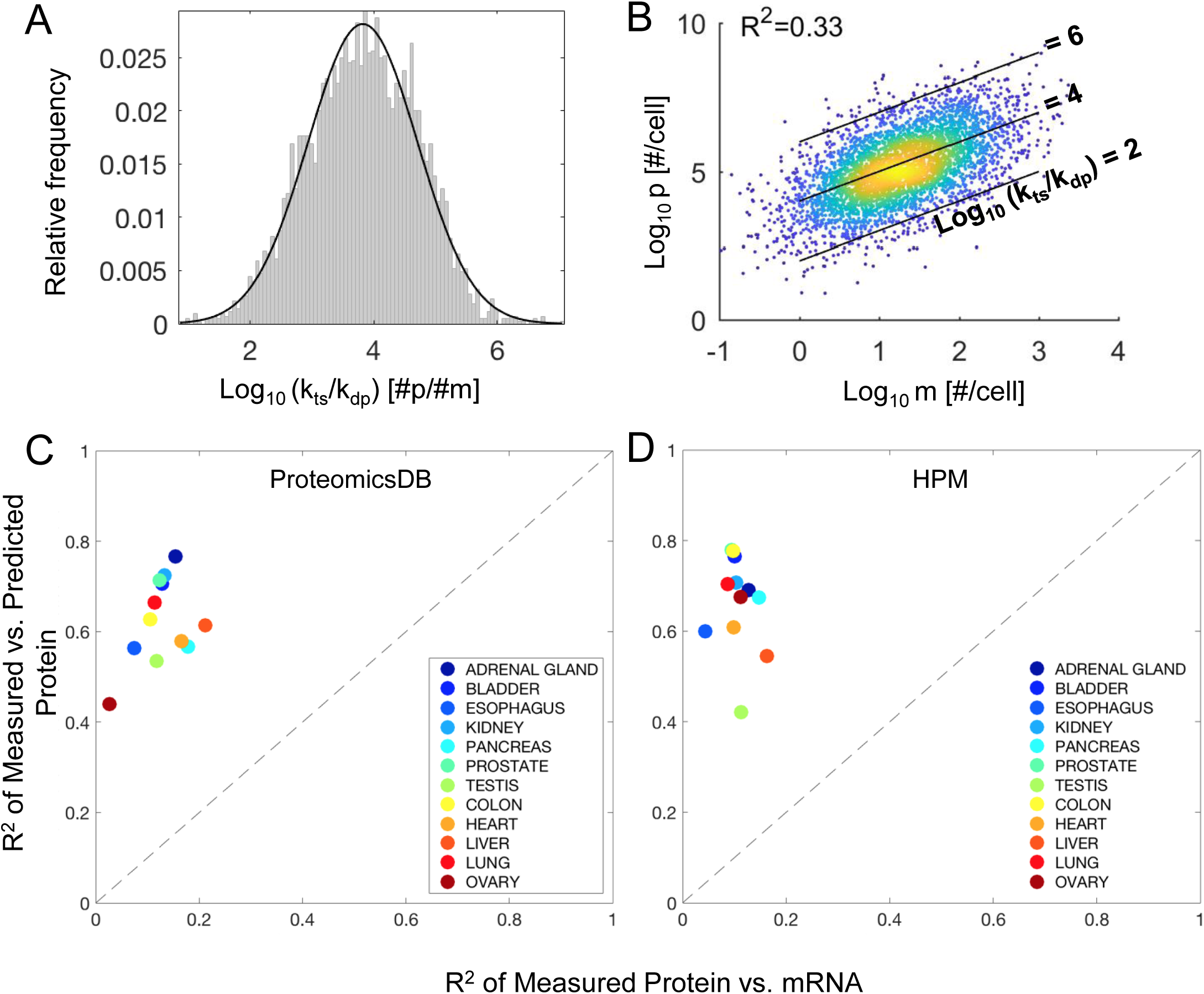
Protein levels correlate poorly with mRNA levels but well with ratio-based predictions. A. First order translation (*k*_*ts*_) and degradation (*k*_*dp*_) rate constants were taken from Schwanhausser et al., and a histogram for the 4,247 genes where both values were available was made for the ratio of the two. The log_10_ values of this ratio are plotted on the x-axis, and their relative frequency is indicated by the gray bars. The data fit well to a log-normal distribution (maximum likelihood), shown as the solid black line, with normal distribution parameters μ=3.82 and σ=0.89. **B.** Both mRNA levels and rate constant ratios were sampled from independent log-normal distributions, and 4,000 protein levels calculated according to Eq. 1. The log_10_ values of the sampled mRNA levels and calculated protein levels are plotted, with color reflecting density of points (blue**→**yellow; low**→**high). Pearson’s R^2^ is calculated from the log_10_ values. **C.** Scatterplot of Pearson’s R^2^ (from log_10_ values) of mRNA vs. measured protein (x-axis) against predicted protein vs. measured protein (y-axis). Marked improvement in R^2^ is seen by all tissues (indicated by color) lying far above the x=y line (black dashes). All protein data were obtained from ProteomicsDB. **D.** A similar analysis in **C** using protein data obtained from Human Proteome Map.

Despite the wide inter-gene rate constant ratio variability, if the ratio for a particular gene were relatively conserved across cell and/or tissue types, then one may be able to improve prediction of protein levels from mRNA levels by using this gene-specific ratio. The public availability of large-scale datasets probing mRNA levels (Genotype Tissue Expression Project-GTEx (Ardlie *et al*, 2015; Melé *et al*, 2015), in units of **r**eads/**k**ilobase of transcript length/**m**illion mapped reads, RPKM), and protein levels from two databases, the Human Proteome Map (HPM) (Kim *et al*, 2014) and ProteomicsDB (Wilhelm *et al*, 2014), in units of spectral counts and iBAQ score, respectively, allowed us to explore this hypothesis. We found 16,708 common unique genes among 12 shared human tissues between these resources (see Methods for details and Table S1). Protein and mRNA levels were measured from several human subjects, but the individual subjects are not matched between the datasets. We calculated the ratio of protein levels to mRNA levels for each gene in each tissue, then took the median ratio across tissues. We used this ratio, along with mRNA data, *m*_i,t_, to predict protein levels for each gene in each tissue, 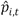, with subscript t denoting tissue. Thus, a single tissue-median ratio, *r*_*i*_,was used across the 12 tissues, but the ratio varied from gene to gene:

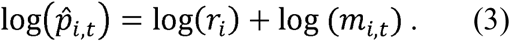

To assess the amount of information contained in these single, tissue-independent but gene-specific ratios, we compared the coefficient of determination (*R*^*2*^) between mRNA and measured protein, to that between measured protein and predicted protein for each tissue (Fig. 1C-D; Supp. Fig. S2). *R*^*2*^ is related to the fraction of variance explained by the independent variable. We found that mRNA variation accounts for roughly 10% of protein level variation (average across tissues of 0.13 for ProteomicsDB and 0.11 for HPM). This is expected and consistent with previous reports (Kosti *et al*, 2016). Using these tissue-independent, gene-specific ratios to predict protein levels increases the *R*^*2*^ from ∼0.10 to ∼0.65 (average across tissues of 0.63 for ProteomicsDB and 0.66 for HPM), increasing the amount of variation explained by ∼55%. This increase is largely consistent across the investigated tissue types, with a slight exception for reproductive tissues (ovary in ProteomicsDB and testis in HPM); however, this trend was not reproduced in both protein databases. Thus, a single gene-specific, but tissue-independent mRNA-to-protein ratio accounts for a significant amount of protein level variation across the human genome. Thus, much of the protein abundance differences across genes are explained by gene-specific translational efficiency and/or protein stability, rather than by mRNA level. Yet, a large amount of variation, ∼35% for the average tissue, remains unexplained. Part of this variation is attributable to experimental error (the % of which is difficult to determine from these data), and also person-to-person and cohort-to-cohort variation reflected by the nature of the databases being comprised of samples from different individuals. However, a significant amount of this remaining variability is likely due to translation or protein stability regulation, which we refer to hereafter as “ratio regulation”.

### Classifying Gene-Specific Predictability Based on mRNA and Protein Level Regulation Across Tissues

Accounting for protein abundance variation across genes does not always imply accurate protein level predictions for a particular gene (Fortelny *et al*, 2017). Although *R*^*2*^ is typically a useful metric for analyzing such prediction tasks, it can be misinterpreted when analyzed on a gene-specific basis. In an extreme example, a protein could be expressed with a constant ratio across 11 tissues, but be radically different in the 12^th^ tissue, leading to poor protein level predictability due to the outlier but a high gene-specific *R*^*2*^ value due to the relative concordance between tissues. Likewise, a relatively low gene-specific *R*^*2*^ value may not imply poor predictive power. For example, there simply could be few ratios available for a given gene due to undetectable expression in several tissues. Alternatively, a gene may be constitutively expressed across tissues at constant mRNA and protein levels, rendering correlation low even though predictability is high.

To help alleviate these gene-specific analysis issues, we calculated mRNA or ratio “variability” *v*_*i*_ for any gene *i* that has four or greater ratio values:

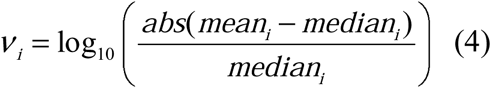

where the mean and median are defined across tissues. A variability of zero or less dictates that the mean is within +/- 1-fold of the median value, implying relative predictability across tissues, and greater than zero, vice versa. In a two-dimensional space (Fig. 2A-C), genes with high mRNA variability and low ratio variability should be the most predictable (Q4). Genes with low mRNA variability and low ratio variability (Q3) are also more predictable; however, the extent of predictability is unclear as both mRNA and protein levels are consistent across tissues, and we have not observed that changes in mRNA levels lead to changes in protein abundances. Genes with high ratio variability (Q2) are of course less predictable, and those with high mRNA and ratio variability (Q1) should be the least predictable. Roughly half the genes have undefined ratios, mainly because protein was not detected in four or more tissues (Fig. 2D-E). Remaining genes were divided into quadrants relatively evenly to analyze their predictability (Fig. 2D-E). Ratio variability for a particular gene did not depend on the number of tissues where ratio was defined (Fig. S3). All genes, their quadrant classification, and their ratio values for each database and tissue are reported in Supplementary Table S2.

**Figure 2.**
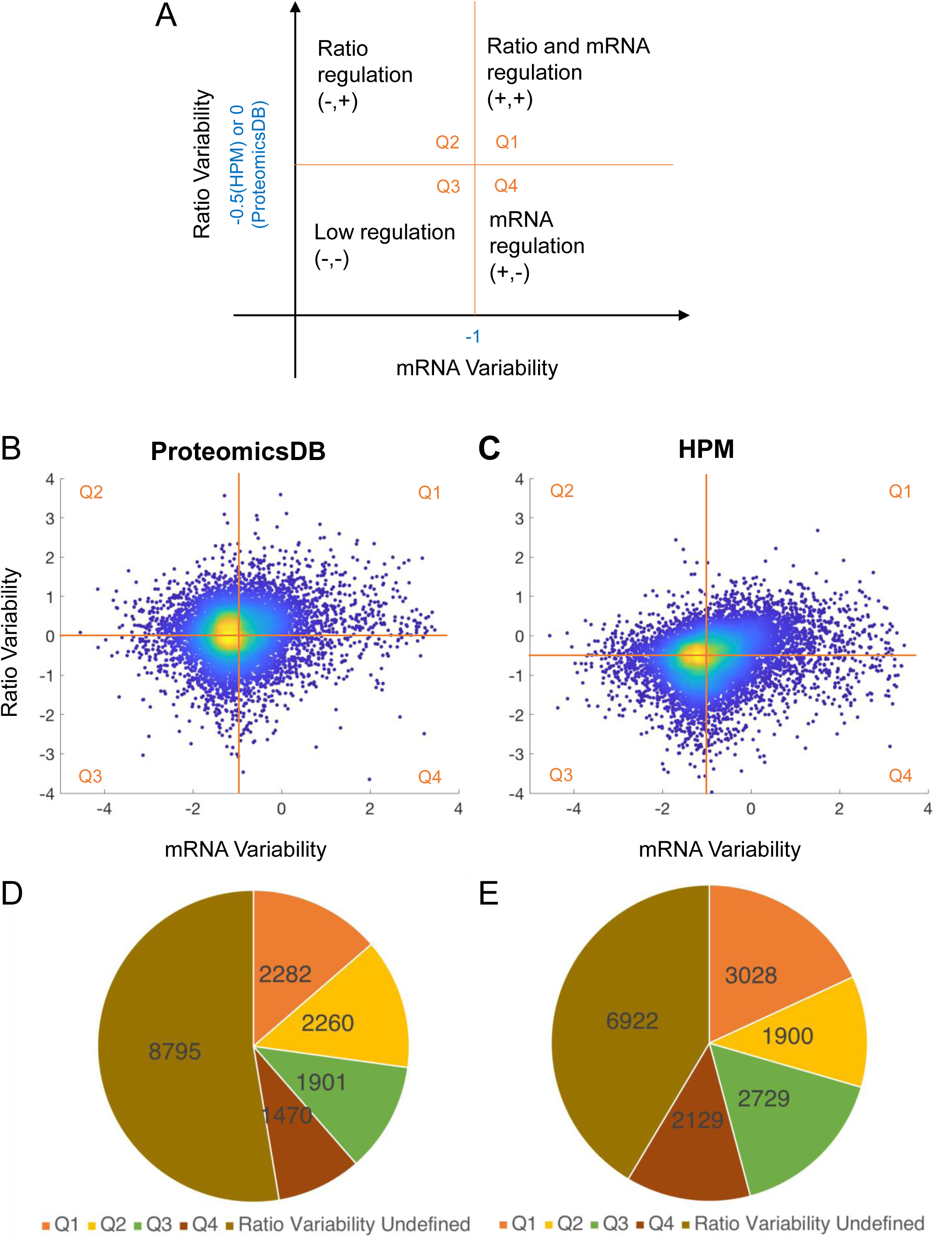
Genes are divided into four quadrants based on their mRNA and ratio variabilities A. Genes are grouped into four quadrants based on their mRNA and ratio variabilities (Eq. 4— log10(abs(mean-median)/median). Orange lines represent the cutoff between quadrants. The cutoff values for each quantity are shown on the axis labels. **B-C.** Ratio variability and mRNA variability for each gene with quadrant cutoffs from ProteomicsDB (B) or HPM (C). Only genes with more than four ratio values were included in this and subsequent analyses. Color indicates density of points. **D-E.** Number of genes in each quadrant for ProteomicsDB (D) or HPM (E).

It is clear from the continuous nature of the variability distributions in Fig. 2B-C that genes at quadrant boundaries have very similar predictability properties. Thus, protein level predictability is likely best understood as a continuum, rather than as a discrete categorical property. Moreover, categorical cutoffs may be tissue and/or cell-type dependent, and also could be shifted depending on the intended application of predicted protein levels and the impact of error on them. Because of the strict cutoff, gene quadrant predictability classifications are not completely overlapping between databases (Table S2); quantitative analysis here employs database-specific definitions which we harmonize later below. Nevertheless, this simple and uniform parsing permitted us to explore hypotheses related to such protein level prediction properties in a broad sense.

### Gene Quadrant Category Strongly Influences Protein Level Predictability

We then asked whether there is any correspondence between *R*^*2*^, across genes for measured vs. predicted protein as above, and quadrant classification. Specifically, we broadly expected *R*^*2*^ values for Q1 and Q2 to be less than those for Q3 and Q4. In support of this hypothesis, in both ProteomicsDB and HPM, we find *R*^*2*^ values to be significantly higher for genes with low ratio variability (Q3 and Q4) as opposed to those with high ratio variability (Q1 and Q2) (Fig. 3A-B). Genes with low ratio variability have ∼75-85% of their protein abundance variance explained by mRNA levels plus a single gene-specific mRNA-to-protein ratio in most tissues and in both databases. Although the amount of protein level variability accounted for by experimental error is difficult to estimate, accounting for 75-85% of protein level variance is approaching what is likely to be close to an upper limit, given the inter-human subject and cohort variability inherent in these studies, in addition to technical variability, although inter-tissue differences in translational and/or post-translational regulation strategies cannot be completely eliminated. There are some tissues for which genes with low ratio variability have unusually low *R*^*2*^, which again include reproductive tissues (particularly Q4), but these data are not consistent across ProteomicsDB (ovary) and HPM (testis), so firm conclusions regarding this are not yet possible. Q4 genes are characterized by high variability in both mRNA levels and ratio across tissues, which includes both mRNA and protein half-life regulation, so it is conceivable that sample integrity issues may be contributing.

**Figure 3.**
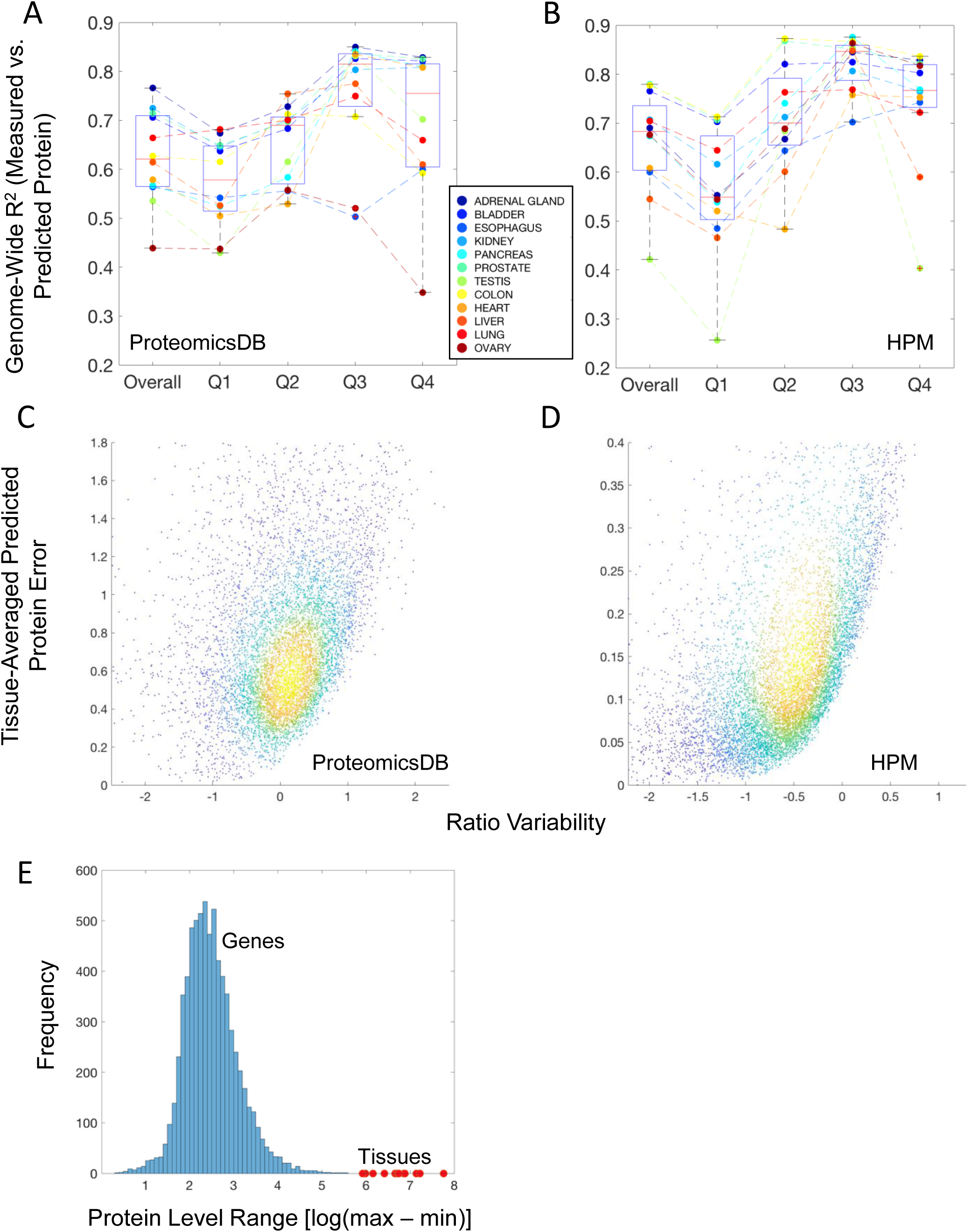
Protein prediction fidelity differs between the four variability quadrants. A-B. The distribution of Pearson’s *R*^*2*^ between predicted and measured protein levels overall and in each quadrant from ProteomicsDB (A) or HPM (B). Points inside each box and whisker indicate a tissue (see color legend). Tissues are connected by dashed lines of the same color. **C-D.** The mean squared error between predicted and measured protein values versus ratio variability for ProteomicsDB (C) or HPM (D). Point density is indicated by the color (blue**→**yellow; low**→**high). The complete data range is in Fig. S4.

Error in predicting protein abundance from mRNA levels plus a single ratio should also increase with ratio variability. We analyzed this error for all genes with a defined ratio, as averaged across all tissues, and find both databases to exhibit this trend (Fig. 3C-D—full scale in Fig. S4). The shape of the prediction error dependence on ratio variability is somewhat different between the two databases, which could be due to the different units of protein abundance reported (ProteomicsDB—iBAQ; HPM—spectral counts), or higher relative ratio variability in ProteomicsDB. Regardless, we concluded that genes with a higher ratio variability are more likely to have larger prediction error across human tissues. Moreover, prediction error is a smoothly varying function of ratio variability, again supporting the notion that protein abundance predictability is best understood as a continuum.

These analyses demonstrate that prediction of protein abundances from mRNA data plus a single tissue-median ratio is greatly improved for genes which are classified here as having low ratio variability (Q3 and Q4). This suggests that gene-specific protein level ranges are intrinsically constrained by a gene-specific ratio; indeed, the range of protein levels for any single gene across tissues is vastly smaller than the range of levels found proteome wide in any tissue (Fig. 3E). Although in these analyses, genes with higher mRNA variability have slightly lower *R*^*2*^, we contend that when extrapolating to other tissues or datasets, such genes would be inherently lower confidence, because we have not observed that changed mRNA levels lead to changed protein levels. Furthermore, these quadrants are broadly defined, and specific protein prediction tasks could benefit from a revised cutoff rationale. For example, a cutoff that varies probabilistically may be beneficial, since prediction error is a continuous function of ratio. If one would like to predict protein abundances in a tissue for which data already exist, one could also use the tissue-specific ratios (Supplementary Table S3), rather than tissue-median ratios as we have done here (Eq. 3). However, units of measurement may become an issue, particularly for the spectral count units in HPM that are more difficult to convert between analyses and are not comparable across genes. In this sense, we stress that the iBAQ units reported by ProteomicsDB are better suited, as they are inherently comparable across genes. Lastly, there are a large number of genes that do not have defined ratio variability because they are not widely expressed across human tissues. Without more characterization both across human cells and tissues, as well as more independent data gathering efforts, prediction of protein abundances for such genes will remain more uncertain, although as noted above, tissue-specific ratios could still be reasonably employed.

### Evaluating Protein Abundance Predictability Classifications with Independent Datasets

How do these protein abundance prediction classifications perform in independent datasets and cell types, particularly when translation rates are likely to be different? To investigate this, we analyzed mRNA and protein abundance data from two serum- and growth-factor starved cell lines: glioma U87 and non-transformed, breast epithelial MCF10A. These two cell lines are clearly disparate from each other, but also, because they were serum- and growth-factor starved, we expect gross changes in translational regulation and capacity as compared to the *in vivo* tissue contexts in the large database resources analyzed above (GTEx, ProteomicsDB and HPM). The cell line protein abundance data are in units of iBAQ score, so we use ProteomicsDB quadrant definitions and tissue-median ratios here.

Correlation between mRNA and protein abundance was expectedly low (*R*^*2*^=0.21 for both lines; Fig. 4A-B). Using the tissue-median ratio from ProteomicsDB to predict protein levels in U87 and MCF10A improves *R*^*2*^ (Fig. 4C-D). In both 10A and U87, genes with high ratio variability in ProteomicsDB (Fig. 4C-D—top row) have significantly lower *R*^*2*^ than genes with low ratio variability (Fig. 4C-D—bottom row), consistent with above analyses. These results demonstrate that ratios defined by tissue-medians in ProteomicsDB can be used to improve predictions of protein abundances in independent datasets, and that prediction is improved much more for low ratio variability genes. Furthermore, the predictions are improved in quite a different biological context: cell lines, both malignant (U87) and non-transformed (MCF10A), as well as in a microenvironment not containing typical serum and growth factor components present in most of the tissues analyzed in ProteomicsDB. We expect that brain- and breast-specific ratios could further improve these predictions, but those data are not yet available in ProteomicsDB to our knowledge.

**Figure 4.**
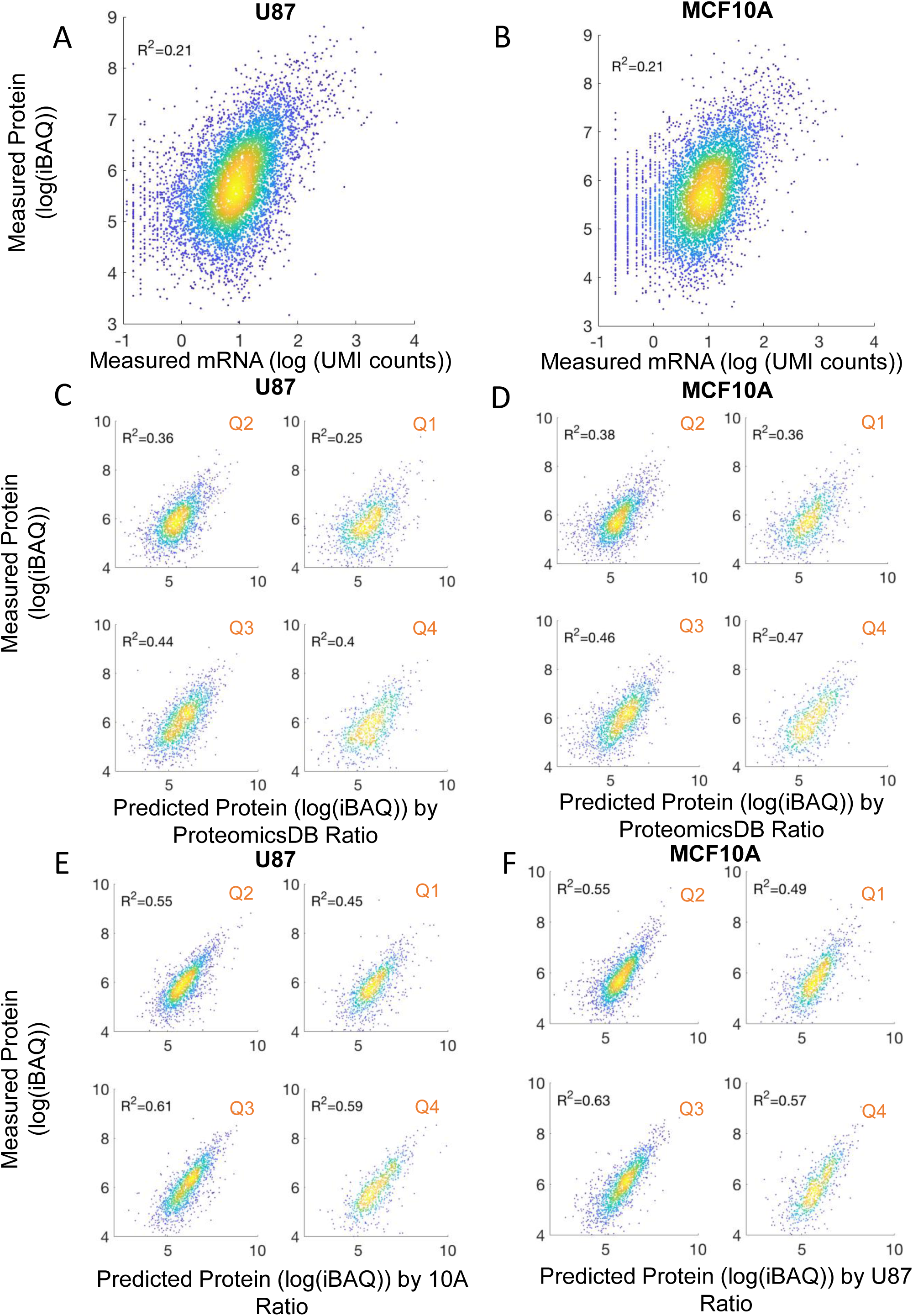
Protein predictions in U87 and MCF10A cell lines. A-B. Protein and mRNA levels of genes for U87 (**A**) and MCF10A (**B**) cell lines. Pearson’s R^2^ is shown in each plot. **C-D.** The measured and predicted protein levels for U87 (**C**) and MCF10A (**D**) cell lines for genes in each of the four quadrants based on ProteomicsDB data. The prediction was given by the gene’s measured mRNA value multiplied by the global tissue-median ratio calculated from ProteomicsDB. Quadrant definitions are indicated and are as in Fig. 2A. **E-F**. The measured and predicted protein levels for U87 (**E**) and MCF10A (**F**) cell lines for genes in each of the four ProteomicsDB-defined quadrants. Here, prediction is based on the gene’s mRNA level in the corresponding cell line multiplied by the ratio from the other cell line. In all panels, ∼5,000-6,000 genes are used depending on data overlap.

We reasoned that using ratios as defined by the serum- and growth-factor starved context could be additionally informative over the ratios given by ProteomicsDB. Therefore, we evaluated how the MCF10A ratios performed for predicting U87 cell protein abundances, and vice versa (Fig. 4E-F). These analyses revealed further significant improvements in *R*^*2*^ in all quadrants, with those genes in Q2 and particularly Q1 remaining reduced compared to other genes. Thus, ratios for prediction that are tuned by microenvironment information may improve protein abundance predictions.

### Enriched Functions of Consensus Genes

Do genes that have more predictable protein abundances have specific biological roles? The quadrant definitions based on ProteomicsDB and HPM are not completely overlapping, mainly because a hard cutoff is used to classify based on the smoothly changing ratio variability. However, we reasoned that genes appearing in the same quadrant from both databases were highly likely to be informative for answering this question. Approximately half of the database-specific genes overlapped, as compared to a quarter by chance (Q1:687, Q2: 1284, Q3: 947, Q4: 1199—Supplementary Table S2). We used Network2Canvas (Tan *et al*, 2013) to identify gene ontology terms that were enriched in biological process and molecular function hierarchies (Fig. 5).

**Figure 5.**
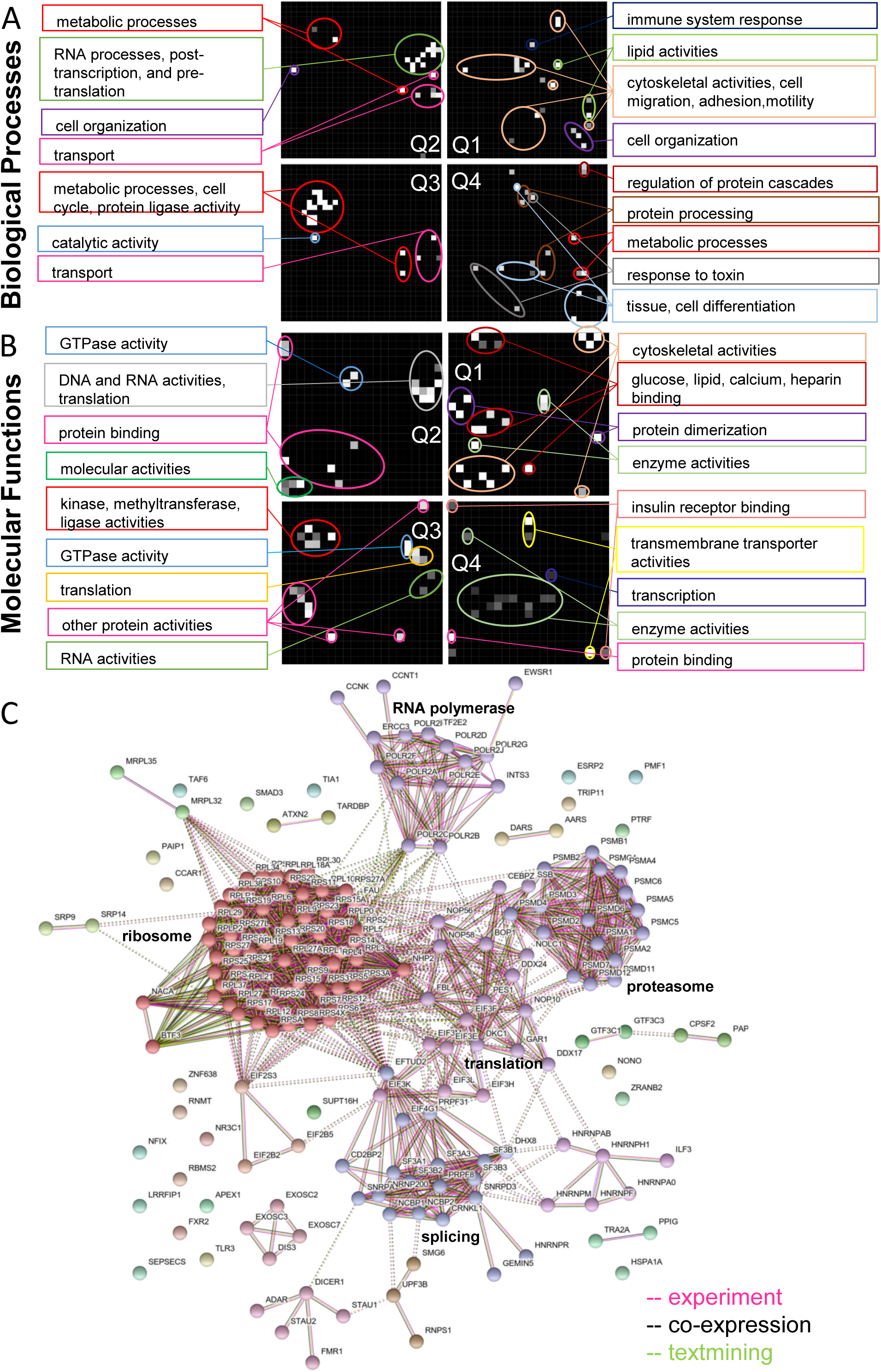
Enrichment canvases show the enriched Gene Ontology terms in each quadrant. A-B. Consensus quadrant genes were taken as those consistently defined in both ProteomicsDB and HPM. Network2Canvas was used to identify the enriched Gene Ontology biological processes (A) and molecular functions (B) terms from consensus genes in each quadrant. Quadrant definitions are as in Fig. 2A. Each square represents each gene ontology term where black squares represent non-enriched terms. Statisical strength is shown by brighter (white) color. Each canvas is annotated with descriptions shown outside the canvas connected to the corresponding squares by color-coded lines. **C.** Network generated from STRING shows relationships between consensus Q2 genes that are involved in pre-translation and post-transcription ontologies. Each node denotes the gene with gene name in the label. The color of the node represents the cluster the gene belongs to. The edges connecting the nodes are color-coded: purple represents experimentally determined, black co-expression, and green text mining.

For more predictable genes with low ratio regulation (bottom quadrants Q3/Q4), metabolic functions, transmembrane transporter activity and tissue development/cell differentiation roles were enriched. Metabolic function genes had low mRNA variability, whereas transmembrane transporter and development/differentiation genes had high mRNA variability. This is consistent with transcriptional regulation of many cell differentiation and developmental circuits (Reik, 2007), as well as cellular stress and xenobiotic response mechanisms via transporter expression (Scotto, 2003). Genes with lower predictability (top quadrants Q1/Q2) are associated with cell adhesion, motility and organization, the immune system, and the cytoskeleton. Post-transcriptional and translation control have been found to be important in a variety of innate, adaptive, chronic, infectious, and autoimmune immunology contexts (Piccirillo *et al*, 2014; Kafasla *et al*, 2014). Interestingly, there is accumulating evidence that the cytoskeleton may itself be a regulator of translation (Kim & Coulombe, 2010), which in turn can locally regulate cell migration through translation control (Liao *et al*; Gu *et al*, 2012).

Surprisingly, genes whose transcripts are constitutively expressed but have less predictable protein levels (Q2) are enriched for DNA/RNA activities, post-transcriptional and translational regulation. The genes associated with these functions and also classified in Q2 by both ProteomicsDB and HPM form recognizable subnetwork clusters related to RNA polymerase, ribosome, proteasome, translation, and RNA splicing (Fig. 5C). This suggests an interesting regulatory mechanism whereby proteins responsible for implementing ratio changes can be themselves quickly up or down regulated to enact the desired changes rapidly in response to signals, or fluctuating conditions. In the case when such network motifs would have an overall negative feedback, they would act as homeostatic controllers to more tightly regulate the abundances of proteins whose levels are important to maintain constant, in a manner that is much more quickly responsive than transcriptional control circuits. We conclude that there are indeed several biological functions that are associated with protein abundance predictability, which gives novel insight to their function and is consistent with growing evidence of ratio regulation-based functions.

## Conclusions

Here we have demonstrated that genes whose proteins have lower ratio variability across human tissues are more likely to have protein abundances that are well predicted using two quantities: an mRNA level and a single gene-specific mRNA-to-protein ratio, taken as the median across human tissues. Ratios from ProteomicsDB were found to apply to independent datasets in cell lines. We provide comprehensive tables as supplementary information to guide others in such prediction tasks. Such prediction is not and will almost certainly never be perfect, but ever-increasing improvements in predictive power will be useful for multiple aspects of biomedical research. Besides such prediction tasks, we found several interesting biological functions for genes depending on how predictable their protein levels are across human tissues, some of which are consistent with accumulating evidence. We uncovered, to our knowledge, a new general network structure whereby genes that control mRNA-to-protein ratios, themselves tend to have their own mRNA-to-protein ratio controlled. This may be a general way for cells and tissues to rapidly enact proteome abundance changes in response to signals, and also for homeostatic control of important protein abundances.

## Acknowledgements

We acknowledge funding to MRB from the NIH Grants R01GM104184, R21CA196418 and U54HG008098 (LINCS Center). Mehdi Bouhaddou and Alan Stern were supported by an NIGMS-funded Integrated Pharmacological Sciences Training Program grant (T32GM062754), DeAnalisa Jones by an NIGMS-funded diversity supplement (R01GM104184-03S1) and an NIH post-baccalaureate research education program (PREP) award (R25GM064118), and Alan Stern by an NIGMS-funded pre-doctoral fellowship (F31GM129985). We thank Mohit Jain and Hong Li for proteomics services, and Evren Azeloglu, Bin (Tina) Hu, Gomathi Jayaraman, and Yuguang Xiong for help with mRNA sequencing. We thank Milo R. Smith for helpful discussions.

## Materials and Methods

### Data Acquisition and Processing

We downloaded data from GTEx (www.gtexportal.org/home/datasets) and HPM (www.humanproteomemap.org/download_hpm_data.php)) on July 20th, 2016. We acquired data from ProtemicsDB through a Request Tool software on July 14th, 2017. All data are in the Supplementary Tables. GTEx data was the median value across human subjects for each tissue type and reported in units of **r**eads **p**er **k**ilo-base of transcript per **m**illion mapped reads (RPKMs). HPM data were similar and in units of spectral counts. ProteomicsDB data were the average value across human subjects (different subjects) in units of log-transformed iBAQ.

The mRNA data from GTEx contained several entries with the same gene ID. For these, we summed the RPKM values for entries that had the same gene ID. We then found matching genes between the two databases which resulted in 16,708 shared unique genes.

The colon, esophagus and heart did not have a direct match between GTEx and HPM. We therefore averaged the data in GTEx from (i) Colon - Sigmoid and Colon – Transverse; (ii) Esophagus - Mucosa, Esophagus - Muscularis, and Esophagus Gastroesophageal Junction; and (iii) Heart - Atrial Appendage and Heart - Left Ventricle. These averaged mRNA level estimates were named colon, esophagus and heart, respectively, and then directly compared to the same. For the ProteomicsDB data, we only included the 16,708 genes shared in the previous two databases for tissues matching the GTEx and HPM databases, resulting in a total of twelve tissues shared between the three databases.

### Cell Culture

MCF10A cells were cultured in DMEM/F12 (Gibco; Cat: 11330032) medium supplemented with 5% (v/v) horse serum (Gibco; Cat: 16050–122), 20ng/mL EGF (PeproTech, Cat: AF-100-15), 0.5 mg/mL hydrocortisone (Sigma, Cat: H-0888), 10μg/mL insulin (Sigma, Cat: I-1882), 100ng/mL cholera toxin (Sigma, Cat: C-8052), and 2mM L-Glutamine (Corning; Cat: 25-005-CI). U87 cells were cultured in DMEM (Gibco, Cat: 10313021) supplemented with 10% (v/v) fetal bovine serum (Gibco, Cat: 10082139). Cells were cultured at 37°C in 5% CO_2_ in a humidified incubator and passaged every 2–3 days with 0.25% trypsin (Corning; Cat: 25-053-CI) to maintain subconfluency. Starvation medium was DMEM/F12 or DMEM medium supplemented with 2mM L-Glutamine. All cell lines were authenticated via STR profiling.

### mRNA Sequencing and Analysis

MCF10A and U87 cells were seeded at ∼30% density (1 million cells / 60mm diameter dish, one dish per biological replicate), and incubated overnight in full growth medium. The next morning, the full growth medium was aspirated, cells were washed once with PBS, and then placed in starvation medium for 24 hours. Total RNA was extracted using TRIzol (Life Technologies, Cat: 15596018) per manufacturer instructions (detailed SOP at www.dtoxs.org; DToxS SOP A– 1.0: Total RNA Isolation). RNA sequencing and analysis was performed as previously described (Xiong *et al*, 2017). Biological triplicates were performed for both cell lines, but one sample in the MCF10A data failed to be sequenced, leaving biological duplicates.

### Proteomics and Analysis

MCF10A and U87 cells were seeded at ∼30% density (5 million cells / 150mm diameter dish, 2 dishes pooled per biological replicate) and incubated overnight in full growth medium. The next morning, the full growth media was aspirated, cells were washed once with PBS, and then placed in starvation medium for 24 hours. Cells were trypsinized for 10 minutes, resuspended in cold (4^°^C) PBS, and spun down at 500g for 5 minutes. Cell pellets were then snap frozen in liquid nitrogen and shipped to the Advanced Proteomics Center at New Jersey School of Medicine to perform mass spectrometry. The cell pellet was subjected to urea lysis buffer composed of 8 M urea, 100 mM TEAB, 1X protease inhibitors (complete, EDTA-free Protease Inhibitor Cocktail from Sigma, prepared according to manufacturer instructions) and 1X phosphatase inhibitors (PhosSTOP from Sigma, prepared according to manufacturer instructions). Protein (50.0 μg total) was resolved by one dimensional SDS-PAGE. The gel lane was divided into 24 equal size bands, followed by in-gel digestion with trypsin (Shevchenko *et al*, 2006). Peptides were fractionated by reverse phase chromatography and analyzed directly by LC/MS/MS on a Q Exactive Orbitrap mass spectrometer (funded in part by NIH grant NS046593, for the support of the Rutgers Neuroproteomics Core Facility). We used Maxquant to analyze the data and calculate iBAQ scores, using a human UniProt protein database as the reference proteome. Final proteins were identified with 1.0% false discovery rate (FDR) at both protein and peptide levels. This resulted in ∼10k unique proteins. Biological triplicates were performed.

**Supplementary Code.** This code (with the associated spreadsheets) can be run in MATLAB to generate all the presented Figures from data (`Figures_Finalized.m`). See `readme.txt`. All scatterplots were made with the use of the function `scatplot` obtained from MATLAB file exchange and was written by Alex Sanchez.

## Supplementary Figures

**Supplementary Figure S1.**
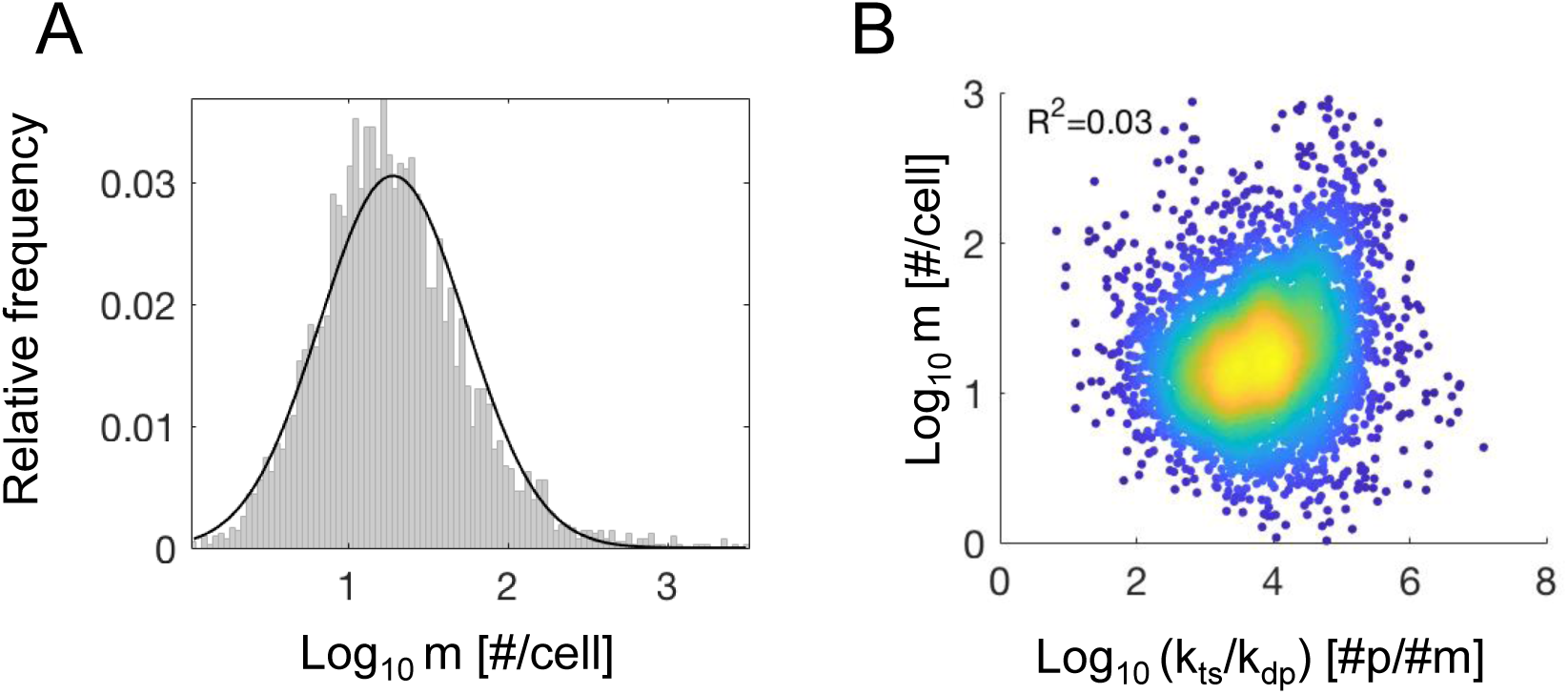
Genomic Variability in mRNA Levels and the Relationship with Rate Constant Ratios. A. mRNA levels were taken from Schwanhauser et al., and a histogram for the 4,309 genes was made. The log_10_ values of this ratio are plotted on the x-axis, and their relative frequency is indicated by the gray bars. The data fit well to a log-normal distribution (maximum likelihood), shown as the solid black line, with normal distribution parameters μ=1.28 and σ=0.46. **B.** The log_10_ mRNA levels and log_10_ rate constant ratios are plotted for the 4,247 genes where ratios exist, with color reflecting density of points (blue**→**yellow; low**→**high). R^2^ is calculated from the log_10_ values, showing a lack of dependency.

**Supplementary Figure S2.**
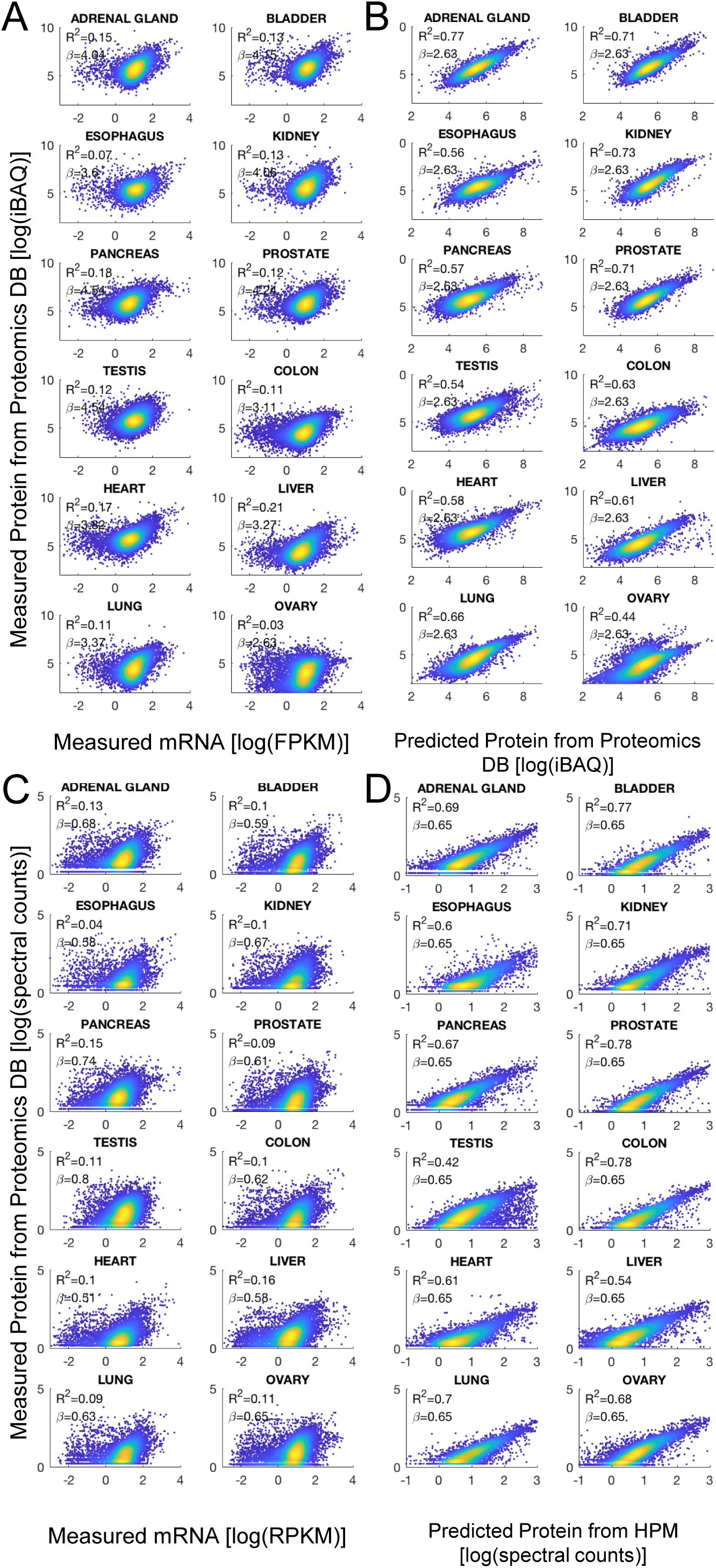
Prediction of protein levels from mRNA levels improve using a single rate constant ratio for all tissues. A,C. The log_10_ of mRNA levels (GTEx) and protein levels (ProteomicsDB for **A** and Human Proteome Map for **C**) for the 16,708 shared genes are plotted for each tissue type, with color indicating density of points (blue**→**yellow; low**→**high). Pearson’s R^2^ is calculated on the log_10_ values as indicated in the top left of each plot. **B,D.** A single estimate of the rate constant ratio for each gene was calculated as the median value across all tissues where the ratio existed (14,803 genes). Predicted protein levels were then calculated by using the tissue-specific mRNA level data, and the median rate constant ratio across all tissues, according to Eq. 1. These predicted protein levels are plotted against measured protein levels in each tissue, with color indicating density of points (blue**→**yellow; low**→**high). Pearson’s R^2^ is calculated on the log_10_ values as indicated in the top left of each plot. Protein data were obtained from ProteomicsDB for **B** and Human Proteome Map for **D.**

**Supplementary Figure S3.**
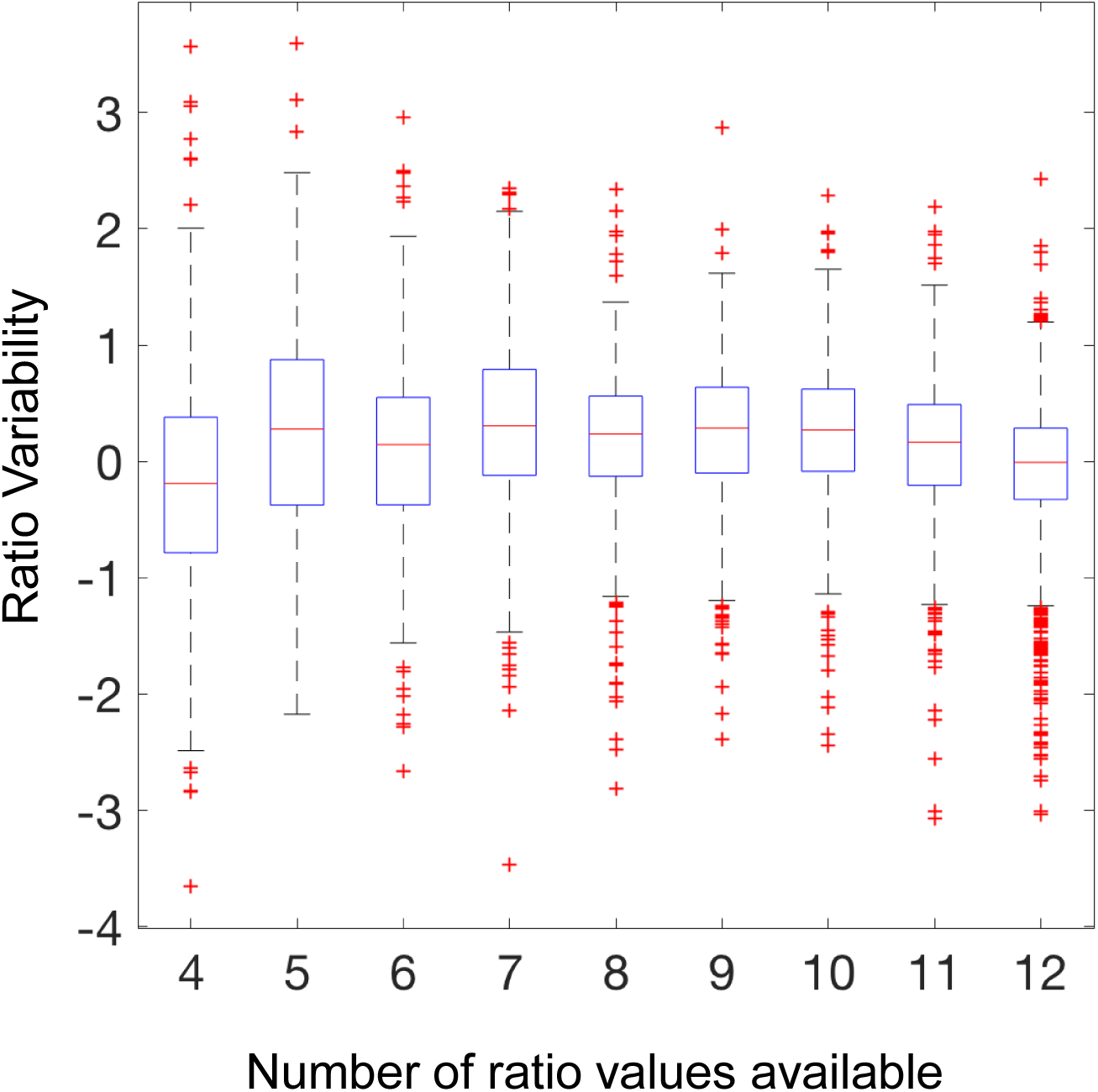
Ratio variability does not depend on the number of values it is calculated from. Each box represents the distribution of ratio variability across genes, conditioned upon the sample size from which ratio was calculated (minimum of four tissues).

**Supplementary Figure S4.**
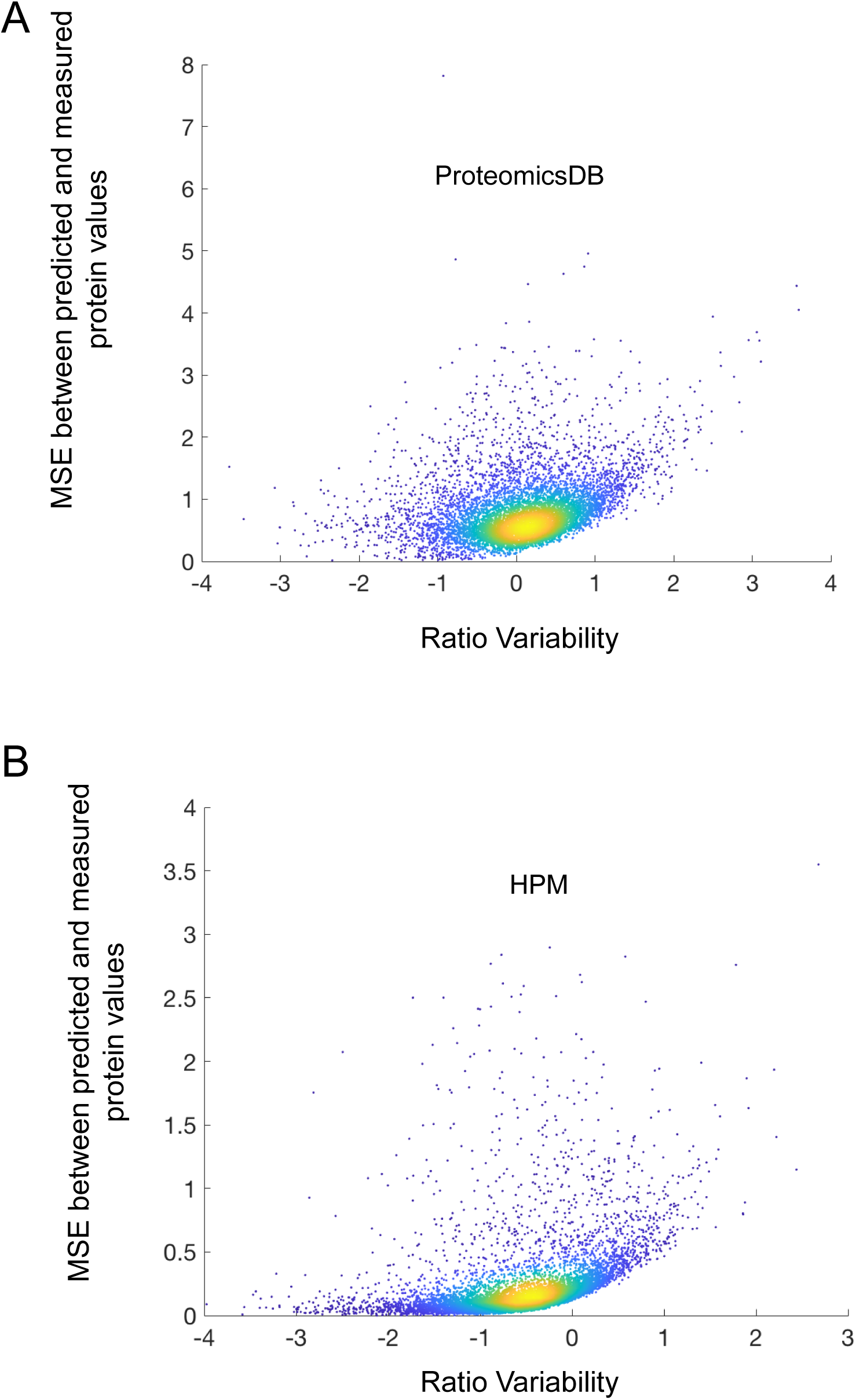
Protein prediction error has a positive relationship with ratio variability. The complete data range for Fig. 3C is shown. MSE: Mean Squared Error.

